# Accurate classification of protein subcellular localization from high throughput microscopy images using deep learning

**DOI:** 10.1101/050757

**Authors:** Tanel Pärnamaa, Leopold Parts

**Affiliations:** Institute of Computer Science, University of Tartu, Tartu, 50409, Estonia; Wellcome Trust Sanger Institute, Hinxton, Cambridgeshire, CB101SA, United Kingdom

## Abstract

High throughput microscopy of many single cells generates high-dimensional data that are far from straightforward to analyze. One important problem is automatically detecting the cellular compartment where a fluorescently tagged protein resides, a task relatively simple for an experienced human, but difficult to automate on a computer. Here, we train an 11-layer neural network on data from mapping thousands of yeast proteins, achieving per cell localization classification accuracy of 91%, and per protein accuracy of 99% on held out images. We confirm that low-level network features correspond to basic image characteristics, while deeper layers separate localization classes. Using this network as a feature calculator, we train standard classifiers that assign proteins to previously unseen compartments after observing only a small number of training examples. Our results are the most accurate subcellular localization classifications to date, and demonstrate the usefulness of deep learning for high throughput microscopy.

---

Microscopy images are a rich, and perhaps underutilized source of high throughput biological data. Endogenous proteins tagged with a fluorescent marker can report quantitative state of living cells, and help annotate gene function by recording spatial and temporal variation in localization or abundance. While biochemical assays of molecule concentrations require large lysed populations for readout, imaging can be performed on single live cells. The acquisition can be automated, producing thousands of micrographs an hour in an arrayed format. These engineering advances have paved way to systematic screening of tagged protein collections^1^, looking for mutant effects on protein abundance^2,3^ and localization^4^, changes in cell^5^ and organelle^6^ morphology, and assigning gene function^7,8^.

Output of a high throughput microscopy screen has to be automatically processed^9^. A typical workflow consists of image normalization, cell segmentation, feature extraction, and statistical analysis; freely available tools exist that make sensible choices for each of these steps^10^^−^^14^. Nevertheless, while the preprocessing stages of normalization and segmentation can be performed in a relatively standardized manner to obtain protein abundances, problem-specific feature extraction and statistical analysis are crucial for subcellular localization mapping. Image analysis pipelines need to carefully calculate more abstract features from raw pixel values, and select most informative ones to obtain numbers that matter in the context of the experiment at hand^15,16^. Defining the correct features can be time consuming and error prone, and default quantities produced by existing software are not necessarily relevant outside the domain for which they were crafted^17,18^.

Deep neural networks^19,20^ have recently become popular for image analysis tasks, as they overcome the feature selection problem. Methods based on deep learning have proved to be most accurate in challenges ranging from object detection^21^ to semantic segmentation^22^ and image captioning^23^, as well as applications to biological domains^24^, from regulatory genomics^25^^−^^27^ to electron microscopy^28,29^. For object identification from photos, these models already outperform humans^21^. Briefly, deep networks process images through consecutive layers of compute units (neurons), which quantify increasingly complex patterns in the data, and are trained to predict observed labels. One of their main appeals is that given a large enough training set, they are able to automatically learn the features most useful for the given classification problem, without a need to design them *a priori*.

Here, we apply the deep learning paradigm to high throughput microscopy data. We present DeepYeast, a neural network trained to classify fluorescent protein subcellular localization in yeast cells. Our network outperforms random forests trained on standard image features for determining the localization patterns, both on single cell and cell population level, and achieves accuracies higher than previously reported. We interpret the internal outputs of the network, and find that neuron layers close to data correspond to low-level image characteristics, while deeper neurons inform of the classification state. The network can be used as a feature extractor, and given the calculated features, separate additional classes using random forests.

## Results

### Deep neural network to classify protein localization in yeast cell images

To perform accurate classification of protein localization in single cells and populations, we created the “DeepYeast” convolutional neural network learned from yeast high throughput microscopy data generated by Chong *et al*.^4^ (Fig. 1a, Supplementary File 1). We used a dataset comprising 90,000 cell images of 1783 proteins localized to exactly one of twelve cellular compartments (Fig. 1b), as measured in two studies^1,4^. Each image records cytoplasmic signal in the red channel, and a fluorescently tagged protein of interest in the green channel. The network consists of 11 layers (eight convolutional layers with rectified linear units, followed by three fully connected layers, Fig. 1c), and a softmax output to assign one of the 12 class labels. DeepYeast’s parameters (over 10,000,000 total) were learned in the Caffe framework^30^, using stochastic gradient descent with momentum (Online Methods).

**Figure 1.**
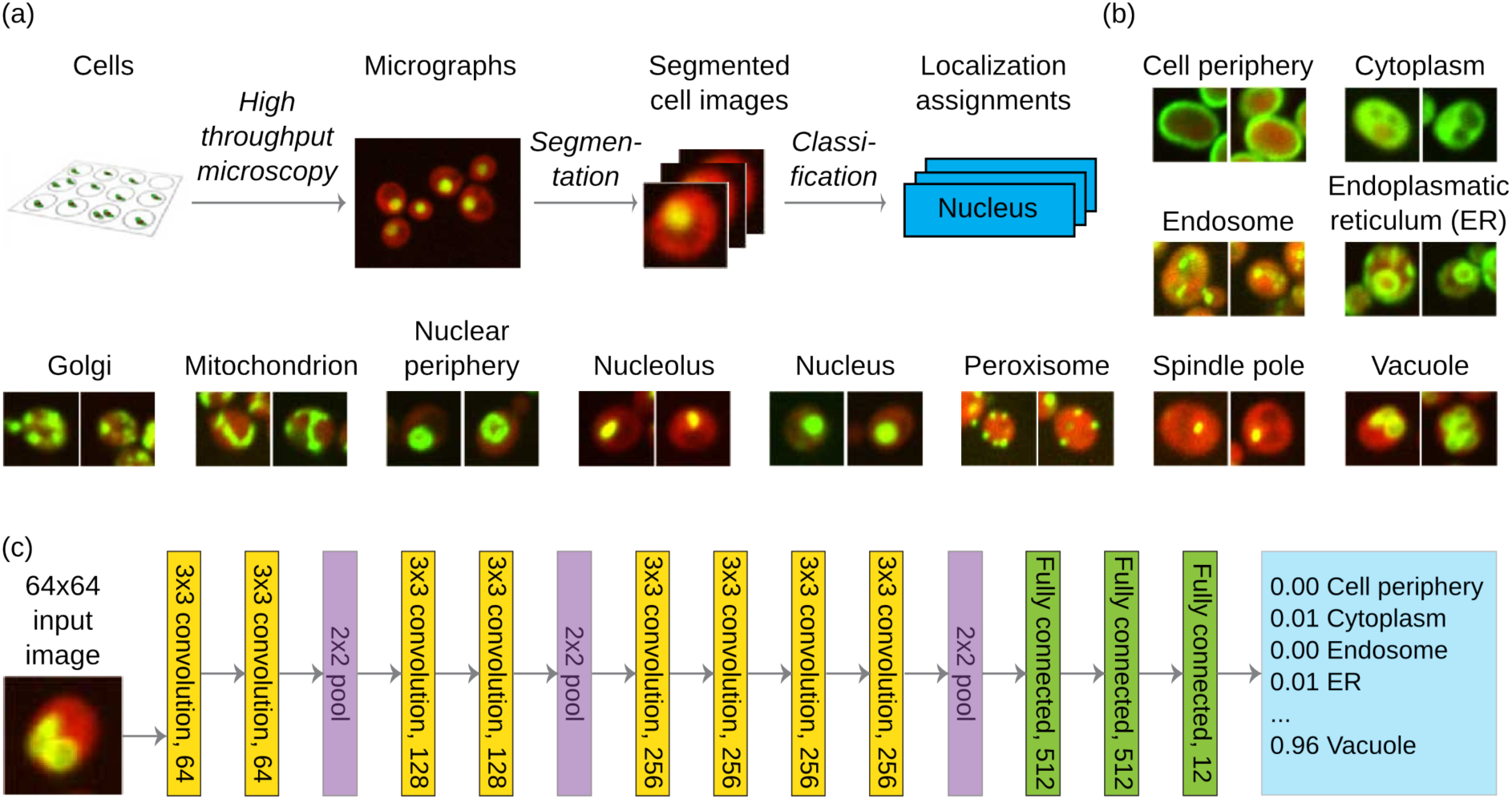
A deep neural network for protein subcellular classification. (a) Outline of the data generation and classification workflow. (b) Example pictures (2 images) from each of the 12 classes (labeled above). Red fluorescence corresponds to a cytosolic marker to denote the cell, and green to the protein of interest. (c) Architecture of the “DeepYeast” convolutional neural network. Eight convolutional layers (yellow) are succeeded by three fully connected ones (green), producing the prediction (blue). All convolutional layers have 3x3 filters with stride 1 (filter size and number of neurons in layer label), and all pooling operations (purple) are over 2x2 non-overlapping areas.

### Accurate classification of protein localization in single cells and populations

We first compared the performance of DeepYeast trained on raw pixel values to random forests^31^ trained on 435 features extracted using CellProfiler^11^ by Chong *et al*.^4^. We fit the models on 72% of the data using a range of parameter settings, picked the one with highest accuracy on another 14% of the data, and quantified its performance on the remaining 14% (Online Methods).

The deep neural network achieved classification accuracy of 87% (10,839/12,500 cells), compared to 75% (9,375/12,500) for random forests (Supplementary File 2,3). DeepYeast outperformed random forests for each class in recall (Fig. 2a) and precision for all compartments except nucleolus (Fig. 2b). The random forest performance is concordant with previous results for single cell classification on the same dataset (70% accuracy^32^), which were obtained using an extended set of classes.

**Figure 2.**
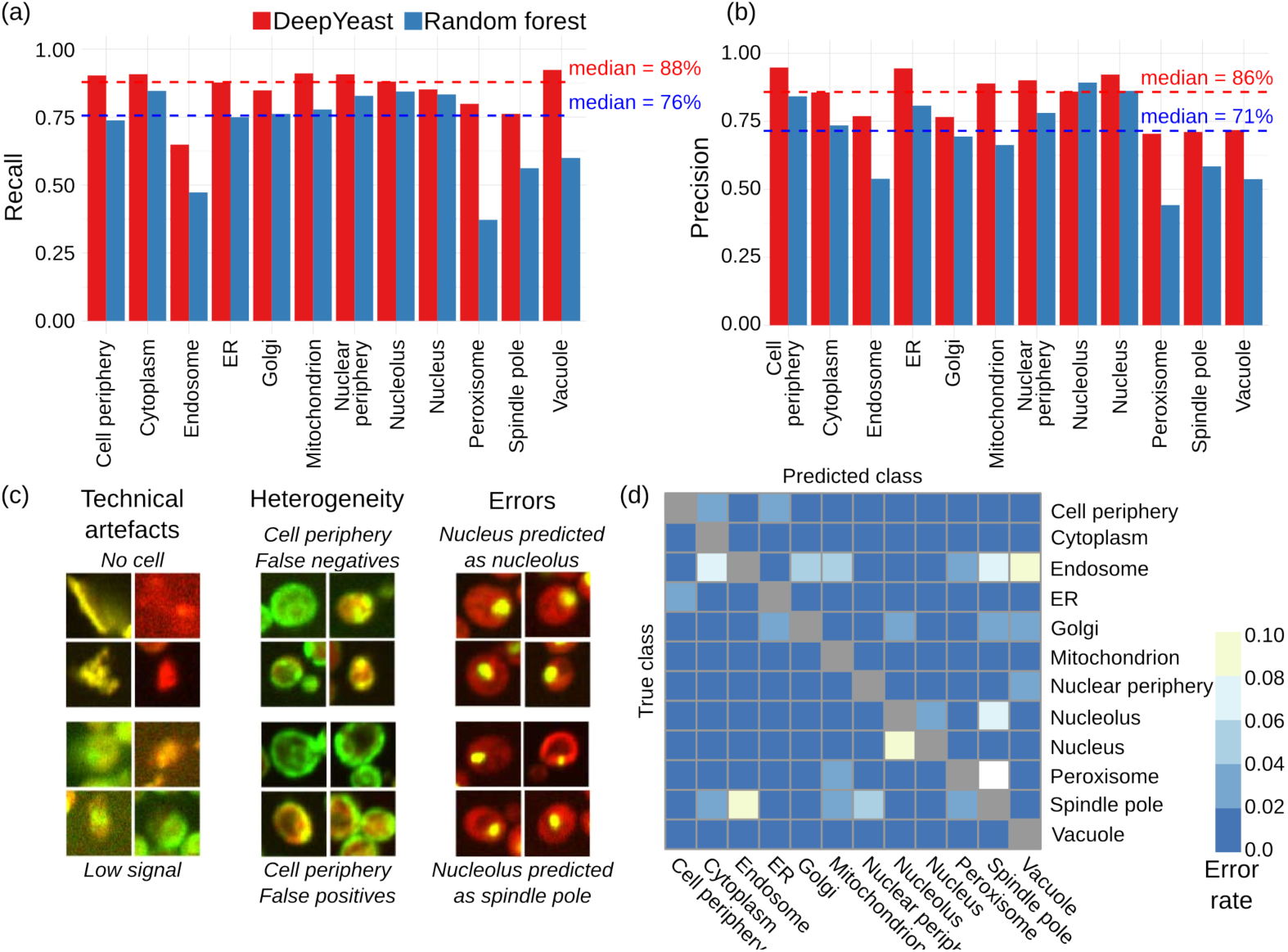
Cellular compartment classification accuracy. (a) DeepYeast outperforms random forests in classification precision. Recall (y-axis) for the twelve subcellular compartments (x-axis) for DeepYeast (red) and random forest (blue) classifiers. The dashed lines denote medians across compartments. (b) Same as (a), but for precision on the y-axis. (c) Example classification mistakes stemming from technical issues (left) due to low signal (bottom left) or no cell (top left), population heterogeneity (middle) resulting in false positives (top middle) and false negatives (bottom middle), as well as frequent model errors (right) of classifying nucleus as nucleolus (top right), or nucleolus as spindle pole (bottom right). (d) Confusion matrix of DeepYeast classification. Error rates from the true (y-axis) to falsely predicted (x-axis) compartments.

Mistakes occurred for each cellular compartment. Some of the errors were due to technical problems with the image, caused by low signal intensity, artefacts, or lack of proper cell (Fig. 2c, left). While our results were generally robust to such noise in the input data, we further trained a classifier to distinguish good quality cell images from the CellProfiler features (Online Methods), and filtered out data deemed to have technical issues, as has been done in previous applications^4^. After removing 1,440 datapoints classified as non-cell (12%), DeepYeast accuracy increased to 91% (10,080/11,060), and random forests to 79% (8,756/11,060). Some remaining errors could be ascribed to labeling mistakes from contamination, or population heterogeneity (Fig. 2c, middle), causing the training data label to be discordant with the observed protein distribution in the cell. In the rest of the cases, DeepYeast classified the protein to the wrong compartment (e.g. Fig. 2c, right).

The most difficult localizations to classify were endosome (recall 65%, 447 correct out of 689), spindle pole (76%, 595/781), peroxisome (80%, 131/164), Golgi (85%, 324/382), and nucleus (85%, 1386/1627). Endosomes, spindle poles, peroxisomes, and Golgi are mainly represented by varying number of puncta, which are not visible in all cells, and may obscure each other, making them difficult to distinguish. Indeed, the most frequent misclassifications (Fig. 2d, Supplementary Fig. 1) were peroxisome to spindle pole (11%; 18 of 164 peroxisome cell images), and endosome to vacuole (8%, 56/689). Another recurring error was designating nucleolar proteins as nuclear (4%, 45/1263), both of which are large round patches. Random forests had additional common mistakes, but the most frequent misclassifications were shared with DeepYeast, reflecting the general difficulty in distinguishing punctate and patch-like patterns in a single cell (Supplementary File 4,5).

So far, we looked at individual cells, and classified the localization pattern of the fluorescent signal. Next, we asked how well we can infer the cellular compartment of a protein from all the cell images acquired for it. We assigned the localization class of each protein as the most probable class according to the posterior probability calculated from aggregating single cell data (Online Methods). Using this combined estimate, we achieved 99% classification accuracy (279/282) on the held out test proteins. Two of the errors were nuclear proteins misclassified as nucleolar, with seven and nine cell images observed, respectively. The remaining error occurred for a protein with a single imaged cell. Thus, further requiring at least ten cells to be recorded for each protein, the accuracy increased to 100% (222/222). Random forests were 95% accurate (269/282) on complete test data, and 96% accurate (214/222) when at least ten cells were measured (Supplementary Fig. 2). As a baseline, Chong *et al*. reported per-protein accuracies of 50-90%^4^ depending on the class on an overlapping dataset, using the Huh *et al*.^1^ annotation as a gold standard. While to our knowledge, human accuracy on these or similar images has not been assessed directly, experts assign compartments to human proteins with over 80% accuracy^32,33^. All these performances are below what we report here, but as the datasets are not identical, direct comparisons should be interpreted with caution.

### Neural network outputs are interpretable

Neural network models are often viewed as black boxes that are difficult to interpret. To gain intuition about DeepYeast features that aid prediction, we analyzed characteristics of learned weights and neuron outputs. We first selected images and image patches that maximize activations of individual neurons, thus matching their weight pattern well (Fig. 3a). The first layers, closest to data, captured general image features, such as sharp edges in the first layer, dots, blobs, and lines in second layer, and more complex boundaries in third and fourth layers. The deeper layers represented combinations of low-level features that started resembling class characteristics, such as punctate patterns, membrane structures, and large patches.

**Figure 3.**
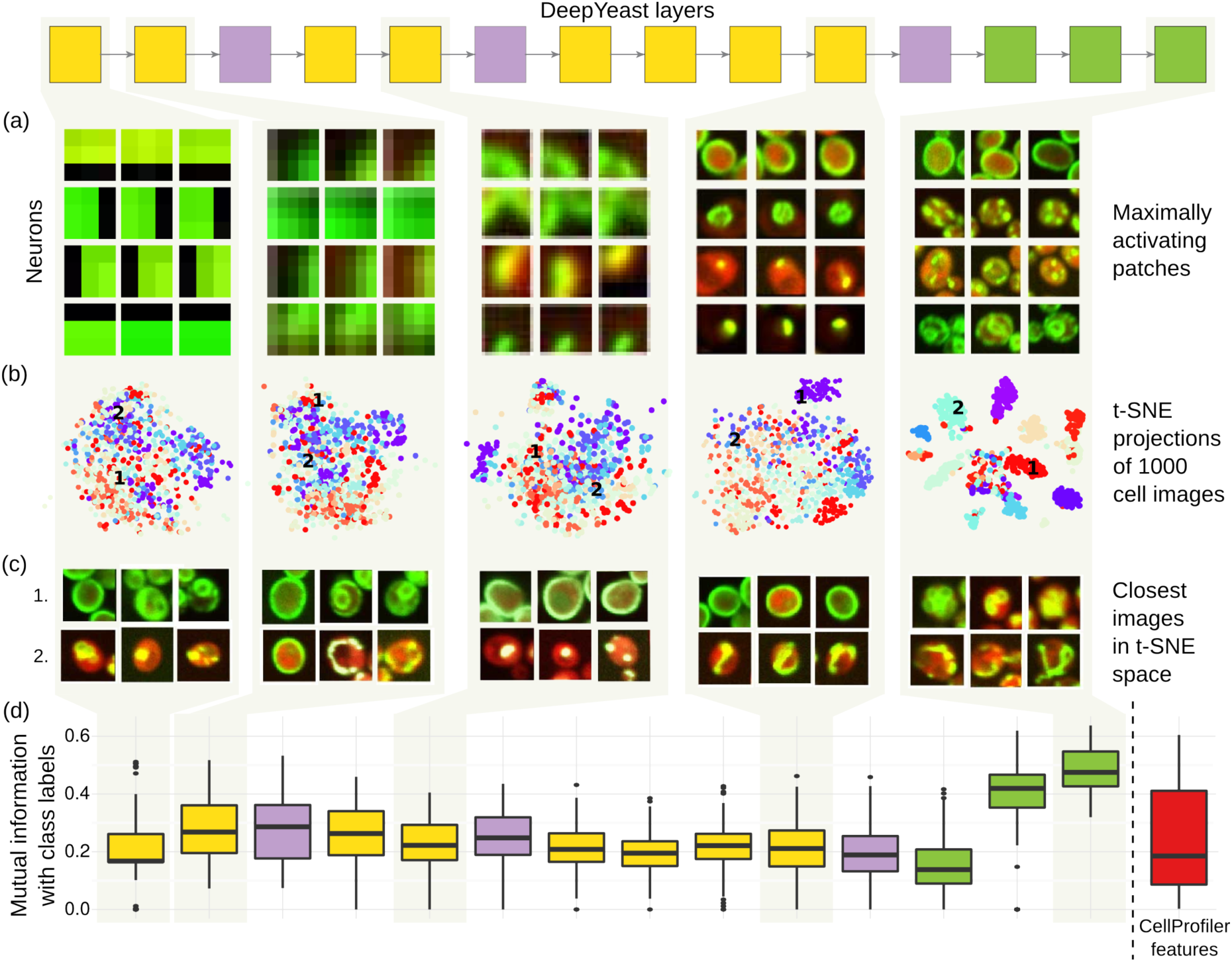
Visualization of the network features at different layers. Interpreting the first, second, fourth, eighth and eleventh layer layers of DeepYeast (box diagram, top, see also Fig. 1c). (a) Image patches that maximize some neuron output. For each of the layers, four neurons (y-axis) and image parts (x-axis) corresponding to a block of pixels that feed into them for maximum activation are shown. (b) 2D visualizations using the t-SNE algorithm^34^. 1,000 random images were fed through the network, hidden layer outputs were extracted, and t-SNE algorithm was used to project the high-dimensional representations into two dimensions. The points are colored based on the true class categories. (c) Three closest images (x-axis) to two chosen points (1 and 2 in Fig. 3b, y-axis) in the two-dimensional t-SNE projection space. (d) Distribution of mutual information (y-axis) between the multinomial class probability and discretized neuron outputs for each layer (left to right), as well as CellProfiler features (rightmost box, red).

Next, we applied t-SNE^34^, a tool to visualize high-dimensional data in two dimensions, on different layers of DeepYeast outputs from 1,000 randomly sampled images, and added compartment information in colors (Fig. 3b, Supplementary Fig. 3). The classes overlap substantially for lower layer outputs, while deeper layers that make use of fully connected network structure increasingly separate the localizations, such that nearby points correspond to the same class (Fig. 3c). We also asked which neuron outputs are correlated to the CellProfiler features and class membership. To do so, we calculated the strongest Pearson correlation coefficient to a CellProfiler feature (as extracted by Chong *et al*.^4^), as well as the largest mutual information with a class label for each unit output. The deep activations informed class labels (Fig. 3d), while shallow ones were more highly correlated to CellProfiler features and Gabor filters (Supplementary Fig. 4).

### DeepYeast can be used as a feature extractor

Classification of new localization classes requires creating new training sets, and if the pattern is rare, obtaining the necessary images is difficult and time-consuming. Further, while applying an existing network to new data can be simple, re-training it requires substantial effort. This motivates repurposing of trained networks as extractors of informative features, which can then be used as inputs to traditional models^35,36^.

We tested whether a network trained on a large amount of data can be used to distill image information that is useful for distinguishing previously unobserved compartments as well. We processed images corresponding to four new challenging classes (actin, bud neck, lipid particle, and microtubule; Fig. 4a) with DeepYeast, and calculated outputs from the first fully connected layer as features. Separation of the classes was evident without additional training (Fig. 4b), indicating that the network extracted informative signal from the data. Next, we trained a random forest classifier on the calculated features, using an increasing number of training images. The classifiers using neural network features outperformed ones using CellProfiler features for small training set sizes (Fig. 4c, Supplementary Fig. 5), and accuracy increased further with additional data. The overall accuracy on these classes remained lower than others due to their punctate pattern, however.

**Figure 4.**
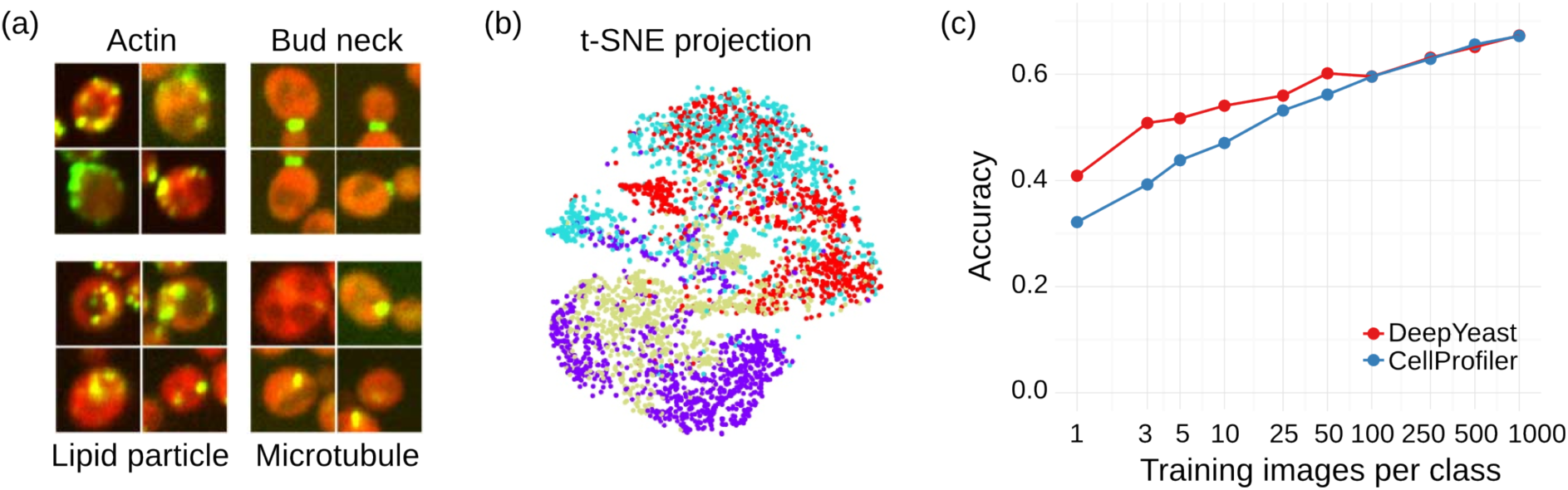
Transfer learning works. (a) Four example images of each of the additional analyzed classes. (b) Applying t-SNE to the network outputs of the additional data (see also Fig. 3b), and coloring the points according to the classes demonstrates separation of new compartments based on features trained for classifying other localizations. (c) Classification accuracy on held-out data (y-axis) for different number of training images (x-axis) for DeepYeast outputs (red) or CellProfiler features (blue) used as inputs to a random forest.

DeepYeast was trained on proteins that predominantly localize to a single compartment. Finally, we confirmed that proteins spread between multiple classes can be accurately inferred as such. We took all proteins manually annotated to belong to nucleus and cytoplasm, and calculated their posterior class probabilities. As expected, cytoplasmic and nuclear classes had high posterior probability, and were the two most probable classes in 21/24 cases, and in the top three for the remaining cases. The per-gene posterior probability of compartment assignment can further be interpreted as the frequency of cells for which the protein resides in the compartment; the model is not forced to make a sharp decision and assign each gene to a single location.

## Discussion

We have demonstrated that DeepYeast, an 11-layer convolutional neural network, can achieve classification accuracy of 91% for individual cells over twelve subcellular localizations, and 100% for proteins when entire cell populations of at least moderate size are considered. Far from being a black box, the internal outputs that DeepYeast produces can be visualized and interpreted in terms of image characteristics. The pre-trained network functions as a feature extractor to successfully distinguish previously unseen classes, and infer mixtures of compartments in a population.

The classification errors mostly occurred between compartments that are also difficult to distinguish by eye. The various numbers of puncta in peroxisome, spindle pole, and endosomes can look like each other, or not be present at all. Nucleus and nucleolus are patches of similar size; when the characteristic crescent shape of the nucleolus is not showing, it is also difficult to distinguish from the nuclear marker. Overall, the single cell accuracy of 91% is approaching the protein compartment assignment performance of previous reports^18,32,33,37^, and the remaining errors are often borderline cases, for which classification is difficult even for trained humans (Supplementary File 4,5). Nevertheless, when at least ten individual cells were measured, the correct cell classifications dominated the errors, and all test proteins were assigned to the right compartment in held out data.

The success of deep neural networks in image analysis relies on architectures that encapsulate a hierarchy of increasingly abstract features relevant for classification, and plentiful training data to learn the model parameters. While first applications used a smaller number of layers^17^, and mostly operated on pre-calculated features^17,18,37,38^, pixel level analyses give good results^39^, especially using the latest training methods^29,32^. Subcellular localization is defined by spatial variation on different length scales, from single small dots to extended thin membranes. Quantification of this covariance structure is thus important for accurate modeling, but deriving the right features for it requires mathematical sophistication and computational crafting^16^. The convolutional layers in the neural network are agnostic to the location of the signal in the image, and take inputs from progressively larger patterns, thus capturing spatial correlations of increasingly wide range in a data-driven manner.

DeepYeast can be reused for other image analysis experiments with the same marker proteins and magnification, or trained further for specific applications. We demonstrated that a pre-trained model can be applied for both classifying previously unseen compartments, and inferring mixtures of localization patterns. The usual classification implementations do not always provide models that are easy to re-use. We envision a repository of networks trained on various bioimage compendia that can be downloaded and employed as out of the box feature calculators, or fine-tuned with additional data to obtain niche-specific results, provided access to the necessary compute. Similar resources already exist in other domains^40^.

While our DeepYeast network outperformed the random forest alternative, and achieved accuracies better than reported before, the direct comparisons must be interpreted with care. We used a clean training set of proteins localized to a single compartment as was done in previous work^4,32^, but as the training data do not match completely, the performance differences may partly be due to the dataset composition. Both Chong *et al*.^4^ and Kraus *et al*.^32^ relied on information from segmented cells for best classification performance; we considered only patches known to contain a cell without pixel-level segmentation information. This circumvents the need for very accurate segmentation pipelines, and indeed, centers of cells can also be derived from additional markers, e.g. histone tags in the nucleus^3^ that are cleanly separated, and therefore much easier to segment than entire cells.

Deep neural networks have proved their value in extracting information from large-scale image data^19,21,41,42^. It would be unreasonable to believe that the same will not be true for high throughput microscopy. Adaptation of the technology will depend on the ease with which it is deployed and shared between researchers; to this end, we have made our trained network freely available. The utility of these approaches will increase with accumulation of publicly shared data, and we expect deep neural networks to prove themselves a powerful class of models for biological image and data analysis.

## Online Methods

### Data

We constructed a large-scale labeled dataset based on high-throughput proteome-scale microscopy images from Chong *et al*^4^. Each image has two channels: a red fluorescent protein (mCherry) with cytosolic localization, thus marking the cell contour, and green fluorescent protein (GFP), tagging an endogenous gene in the 3’ end, and characterizing the abundance and localization of the protein. For ~70% of the yeast proteome, the protein subcellular localization has been manually assigned (Huh *et al*.^1^). However, our data were acquired in a somewhat different genetic background and experimental setting, and labeling the images by eye can be error-prone. To obtain high confidence training examples, we therefore used images where Huh *et al*.^1^ and Chong *et al*.^4^ annotations agree. Our final data set comprised 7,132 microscopy images from 12 classes (cell periphery, cytoplasm, endosome, endoplasmatic reticulum, golgi, mitochondrion, nuclear periphery, nucleolus, nucleus, peroxisome, spindle pole, and vacuole) that were split into training, validation and test sets. Furthermore, segmentations from Chong *et al*.^4^ were used to crop whole images into 64 x 64 pixel patches centered on the cell midpoint, resulting 65,000 examples for training, 12,500 for validation and 12,500 for testing.

### Convolutional neural network

We trained a deep convolutional neural network that has 11 layers (8 convolutional and 3 fully connected) with learnable weights (Fig. 1c). We used 3x3 patterns with step size (stride) 1 for convolutional layers, 2x2 aggregation regions with step size 2 for pooling layers, and rectified linear unit nonlinearities for the activation function. The number of units in the convolutional layers was 64, 64, 128, 128, 256, 256, 256, and 256, and in the fully connected layers 512, 512, and 12. We initialized the weights using Glorot-normal initialization technique^43^, and used batch normalization^44^ after each convolutional or fully connected layer, but before activation functions. For each image, per-pixel training set mean was subtracted before use. Cross-entropy loss was minimized using stochastic gradient descent with momentum of 0.9, initial learning rate of 0.1 and a mini-batch size of 100. Learning rate was divided by 2 after every 16,250 iterations (25 epochs). To reduce overfitting, we used weight decay of 0.0005, and dropout with rate of 0.5 for the first two fully connected layers. The models were trained for 195,000 iterations (300 epochs over full training data), and based on validation loss, the model at iteration 130,000 was chosen for all experiments. The training took three days on an NVidia Tesla K20m graphical processing unit. We also tested networks of depth 5, 9, and 13, all of which performed worse on the validation dataset.

### Random forest

For comparison, we trained a random forest classifier on features from Chong *et al*^4^ that were extracted using a CellProfiler^11^ pipeline. In total, there are 435 different features consisting of intensity, geometric and texture measurements on different scales, such as Haralick texture features^45^, Gabor^46^ and Zernike^47^ filters. We performed a grid search to select the number of trees to grow (50, 100, 250, 500, or 1000), number of features to randomly sample at each split (10, 25, 50, 75, 100, 125, 150, 175, 200, 250, or 300), and minimum size of terminal nodes (1, 2, 5, 10, or 50). Based on validation set performance, we chose 500, 100, and 1 for these hyperparameters, respectively. The final performance was evaluated on the same test dataset as the neural network.

### Protein-level classification

For both random forest and DeepYeast, we modeled the protein localization in one cell as a multinomial distribution with uninformative Dirichlet prior for the protein, and calculated the Dirichlet posterior for the protein from observations of individual cells. We used the maximum *a posteriori* estimate for protein localization. Intuitively, this approach corresponds to softly counting the number of cells assigned with each compartment, and picking the compartment with the maximum count.

### Determining good quality cells

To remove a prominent source of misclassifications, we trained a random forest to discriminate between cells and non-cells (e.g. inappropriately segmented regions, empty areas, and imaging artifacts) based on the CellProfiler features. For each of the twelve categories, we randomly sampled without replacement 100 examples from the validation set images that were correctly classified by both DeepYeast and random forest, and labeled them as good quality cells. In addition, we inspected the validation set, and manually picked 118 non-cells, resulting in a total of 1,200 cell, and 118 non-cell images. We performed 10-fold cross-validation to choose the number of features to randomly sample at each split (2, 110, 218, 326, or 435), and whether to downsample good quality images at every bootstrap sample. Based on cross-validation performance, the final model used 100 trees, 110 features at each split, no downsampling, and achieved accuracy of 96.7%.

### Transfer learning

To assess the generality of DeepYeast features learned in the classification task, we constructed a new data set from classes not present in the training data. The four new categories (actin, bud neck, lipid particle, microtubule) each contained 1,000 cell images for training, 500 for validation and 1,000 for testing. We fed the data into DeepYeast, and extracted the outputs of the first fully connected layer as features. We subsampled random data sets of different sizes (1, 3, 5, 10, 25, 50, 100, 250, 500) from training data, fit a random forest classifier to the corresponding DeepYeast and CellProfiler features, picked the best performing model on validation data, and evaluated the final performance on testing data for every data set size. We also employed L1-regularized logistic regression models, which did not perform better than random forests for this task.

### t-SNE visualizations

We picked 1,000 cells at random across all classes, processed them with the DeepYeast network, and applied t-SNE^34^ with default parameters to the neuron outputs at the different layers.

### Data availability

The data used in this study were described in Chong *et al*.^4^, and stored in the Cyclops database presented in Koh *et al*.^48^

## End Notes

### Author contributions

TP trained the models and analysed data. TP and LP conceived and designed the analysis approaches, and wrote the manuscript.

## Acknowledgements

We thank Helena Friesen for consultations on yeast images, Yolanda Chong, Judice Koh, and Oren Kraus for help with accessing the data, and Martin Hemberg, Jared Simpson, Felicity Allen, Oliver Stegle, and Christof Angermüller for comments on the text.

